# NEK kinase TcRDK2 controls differentiation, host-cell infection, and remodeling of the translation initiation machinery in *Trypanosoma cruzi*

**DOI:** 10.64898/2026.06.07.730777

**Authors:** Juliana N. Roson, Jessica Huckleberry, Noelia Lander, Miguel A. Chiurillo

## Abstract

*Trypanosoma cruzi*, the etiologic agent of Chagas disease, alternates between replicative epimastigotes and amastigotes and non-dividing, mammal-infective metacyclic and bloodstream trypomastigotes. Protein phosphorylation is a major regulatory mechanism in trypanosomatids, whose kinomes reveal an expanded family of NIMA-related kinases (NEKs). Here, we investigated the role of *T. cruzi* RDK2 (Repressor of Differentiation Kinase 2), a conserved NEK that carries a C-terminal pleckstrin homology (PH) domain. Endogenous gene tagging showed that TcRDK2 is expressed in all major life-cycle stages and displays a cytoplasmic distribution. CRISPR/Cas9-mediated knockout of *TcRDK2* did not markedly alter epimastigote growth in rich medium but caused a significant accumulation of cells with abnormal nuclear/kinetoplast configurations, consistent with defects in kinetoplast segregation and cytokinesis; *TcRDK2*-null parasites also showed reduced *in vitro* metacyclogenesis and failed to establish efficient infections in human fibroblasts. To probe gain-of-function effects, we generated tetracycline-inducible overexpression lines for full-length TcRDK2 (RDK2^WT^), a PH-deleted variant (RDK2^ΔPH^), and a catalytic-dead mutant (RDK2^K70A^). Overexpression of RDK2^WT^ or RDK2^ΔPH^ decreased epimastigote growth, enhanced metacyclogenesis, and strongly impaired host-cell invasion and intracellular amastigote proliferation, with more pronounced phenotypes for RDK2^ΔPH^, suggesting that the PH domain normally restrains TcRDK2 activity in vivo. Phosphoproteomic profiling of RDK2^WT^-overexpressing epimastigotes identified candidate TcRDK2 substrates and pathways, including translation initiation and cytoskeletal regulation. Together, these data identify TcRDK2 as a NEK kinase that coordinates kinetoplast replication/segregation, metacyclogenesis, and host-cell infection in *T. cruzi* and support TcRDK2 as a promising, kinetoplastid-specific therapeutic target for Chagas disease.

**IMPORTANCE:** Chagas disease, caused by the parasite *Trypanosoma cruzi*, remains a major health problem with limited treatment options. To persist in both insect vectors and mammalian hosts, the parasite must precisely coordinate cell division, differentiation into infectious forms, and survival inside host cells. Protein kinases are central regulators of these processes and attractive drug targets, yet many remain poorly understood in *T. cruzi*. In this study, we investigate RDK2, a member of the NIMA-related kinase family. Using gene knockout, inducible overexpression, and global analysis of phosphorylated proteins, we show that RDK2 is required for accurate segregation of mitochondrial DNA, efficient formation of infective insect-stage forms, and successful infection and replication in human cells. These findings identify RDK2 as a key regulator that links parasite cell division to infectivity and highlight it as a promising, parasite-specific candidate for future drug development against Chagas disease.

## INTRODUCTION

Chagas disease, caused by the protozoan parasite *Trypanosoma cruzi*, remains the most prevalent parasitic disease in the Americas and has been recently recognized as a low-endemicity infection in the United States, with locally acquired and imported cases increasingly documented (1, 2). Approximately 8 million people are infected worldwide, with tens of thousands of new cases and deaths every year, and a large population at risk of developing chronic cardiomyopathy or digestive megasyndromes (3, 4). Current chemotherapy relies almost exclusively on benznidazole and nifurtimox, drugs developed more than five decades ago that show limited efficacy in the chronic phase of the disease and are frequently associated with severe adverse effects (5, 6). The absence of vaccines and the shortcomings of existing treatments underscore the urgent need to identify novel, parasite-specific targets that can be exploited for safer and more effective chemotherapeutic strategies against Chagas disease.

*T. cruzi* is a kinetoplastid protozoan with a complex digenetic life cycle (7). The parasite alternates between the insect vector and the mammalian host, cycling through four major developmental forms: epimastigotes and metacyclic trypomastigotes in the triatomine gut, and cell-derived trypomastigotes and intracellular amastigotes in mammalian tissues. Among these, the epimastigote and amastigote stages are proliferative, whereas the metacyclic and bloodstream trypomastigotes are non-dividing but infective forms that mediate transmission and establishment of infection. As *T. cruzi* progresses through its life cycle and transitions between these stages, it encounters drastic changes in its environment (including temperature, pH, nutrient availability, and host-derived stresses) to which the parasite must rapidly sense and respond through tightly regulated molecular and cellular adaptations (8, 9).

Protein kinases are central regulators of virtually all aspects of eukaryotic cell biology. By catalyzing reversible phosphorylation of serine, threonine, or tyrosine residues, they modulate enzyme activity, protein-protein interactions, subcellular localization, and protein stability, thereby organizing signaling networks that control cell-cycle progression, differentiation, metabolism, environmental sensing, and stress responses (10, 11). In trypanosomatids, protein phosphorylation has emerged as a dominant mode of post-transcriptional regulation, compensating for the limited transcriptional control resulting from polycistronic transcription and trans-splicing (12, 13). System-level kinome analyses have revealed that *T. cruzi*, *Trypanosoma brucei*, and *Leishmania spp*. possess a complex and highly divergent repertoire of eukaryotic protein kinases (ePKs) and atypical protein kinases (aPKs), many of which have no close human orthologs and are predicted to be essential for parasite survival (14–16). These features make trypanosomatid kinases attractive targets for selective drug development (17, 18).

Within *T. cruzi*, functional studies using reverse genetics and chemical biology have implicated protein kinases in the regulation of cell-cycle progression, kinetoplast replication and segregation, cytokinesis, environmental sensing, parasite viability, infectivity and stage differentiation, and have highlighted that several kinases are essential for epimastigote proliferation and metacyclogenesis, as well as for intracellular replication and host-cell invasion (19–23). We recently developed and applied CRISPR/Cas9-based genome editing in *T. cruzi* to interrogate the function of an essential protein kinase. Using knockout and analog-sensitive mutant cell lines, we showed that the AGC Essential Kinase 1 (AEK1) is required for epimastigote proliferation, host-cell invasion, and intracellular replication, and that its acute inhibition reveals a critical role as a regulator of cytokinesis in this parasite (20). Taken together, this and related studies demonstrated that targeted genetic manipulation of kinases is a powerful approach to dissect kinase-centered signaling networks in *T. cruzi* and other trypanosomatids and to prioritize promising drug targets (22, 24, 25).

Among the trypanosomatid kinases implicated in developmental control, two genes were identified through kinome-wide RNAi screening in *T. brucei* and named Repressor of Differentiation Kinases 1 and 2 (*RDK1* and *RDK2*), because they are protein kinases that in that parasite act as negative regulators of the differentiation process from bloodstream-form to procyclic-form (14). RDK1 is classified as an STE11-like kinase, whereas RDK2 is a member of the NIMA-related kinase (NEK) family of serine/threonine protein kinases, a group widely associated with cell-cycle progression and cytoskeletal dynamics in eukaryotes (26–28). In *T. brucei*, RDK2 acts as a negative regulator of differentiation; its depletion triggers inappropriate expression of procyclic markers, whereas overexpression suppresses differentiation under permissive conditions (14). In *Leishmania mexicana*, attempts to generate RDK2 knockout mutants in procyclic promastigotes were unsuccessful, and *LmRDK2* was therefore considered likely to be essential (29). Despite these findings, that highlight the importance of RDK2 in trypanosomatids, the role of this NEK-family kinase in *T. cruzi* biology has not yet been explored.

Here, we apply a kinase-focused strategy to study the *T. cruzi* ortholog of *RDK2*. Given the established role of the NEK-family kinase RDK2 in differentiation control in *T. brucei* and its reported essentiality in *Leishmania* promastigotes, we hypothesized that TcRDK2 might act at the interface between cell-cycle regulation, kinetoplast segregation, and developmental transitions during the *T. cruzi* life cycle. To test this hypothesis, we used CRISPR/Cas9 technology to generate multiple genetically modified cell lines, including a *TcRDK2* knockout and parasites with endogenously tagged *TcRDK2*. In addition, we generated tetracycline-inducible *TcRDK2* overexpression cell lines using a conditional expression system. Using this toolkit, we provide evidence that TcRDK2 participates in the control of epimastigote proliferation, kinetoplast division, and the capacity *of T. cruzi* to infect and replicate within mammalian host cells. By placing TcRDK2 within the broader framework of NEK-family kinases and *T. cruzi* kinase biology, our work reinforces the view that parasite protein kinases could represent a rich reservoir of novel, parasite-specific drug targets for the treatment of Chagas disease.

## RESULTS

### Expression and localization of the TcRDK2

An *RDK2* ortholog was found in the *T. cruzi* YC6 genome (gene ID: TcYC6_0103940), where it is annotated as a *putative serine/threonine-protein kinase a*, corresponding to a single-copy gene of 1,330 nucleotides encoding a 440 amino acids protein of ∼50 kDa. *RDK2* orthologs are conserved across all kinetoplastid species whose genomes are currently available in public databases, ranging from free-living organisms to mammalian pathogens. In this context, *TcRDK2* exhibits 63% to 79.5% amino acid identity and 88.2% to 97.5% similarity to orthologous from *L. mexicana* and *T. brucei*, respectively. The AlphaFold 3 structural predictions (Fig. S1) show that like most of NEK family members in eukaryotic organisms, the TcRDK2 kinase domain adopts a bilobed fold composed of an N-terminal lobe (N-lobe) and a C-terminal lobe (C-lobe). Key motifs of NEKs present in the catalytic domain of TcRDK2 include a conserved HRD (His165/Arg166/Asp167) motif for catalytic activity, an activation loop residue (Thr207) for phosphorylation, and the DFG (Asp185/Phe186/Gly187) motif involved in ATP (26). In addition, TcRDK2 possess a C-terminal PH domain, a domain architecture that is present in several trypanosomatid NEKs but has not been reported in NEKs from other organisms (15). To determine TcRDK2 cellular localization, we generated an endogenously C-terminal tagged mutant cell line by CRISPR/Cas9 (*TcRDK2*-3xc-Myc) (Fig. 1A). *TcRDK2* tagging was confirmed by PCR (Fig. 1B) and by western blotting using anti c-Myc antibodies (Fig. 1C). Using this cell line, we established the subcellular localization of TcRDK2 in the four developmental stages maintained in laboratory conditions, epimastigote, metacyclic trypomastigote, cell-derived trypomastigote and amastigote. As shown by immunofluorescence assays (IFA), TcRDK2-3xc-Myc displays a cytosolic punctate distribution in all developmental stages (Figure 1D). These observations indicate that TcRDK2 is constitutively expressed throughout the life cycle of *T. cruzi*. To confirm this localization pattern, we generated a mutant cell line in which TcRDK2 was endogenously tagged at its N terminus (Fig. S2A). 3xc-Myc-*TcRDK2* tagging was confirmed by PCR (Fig. S2B) and by western blotting using anti-c-Myc antibodies (Fig. S2C). IFA revealed that TcRDK2 displays the same localization pattern as the C-terminally tagged protein in *T. cruzi* epimastigotes (Fig. S2D).

**Figure 1.**
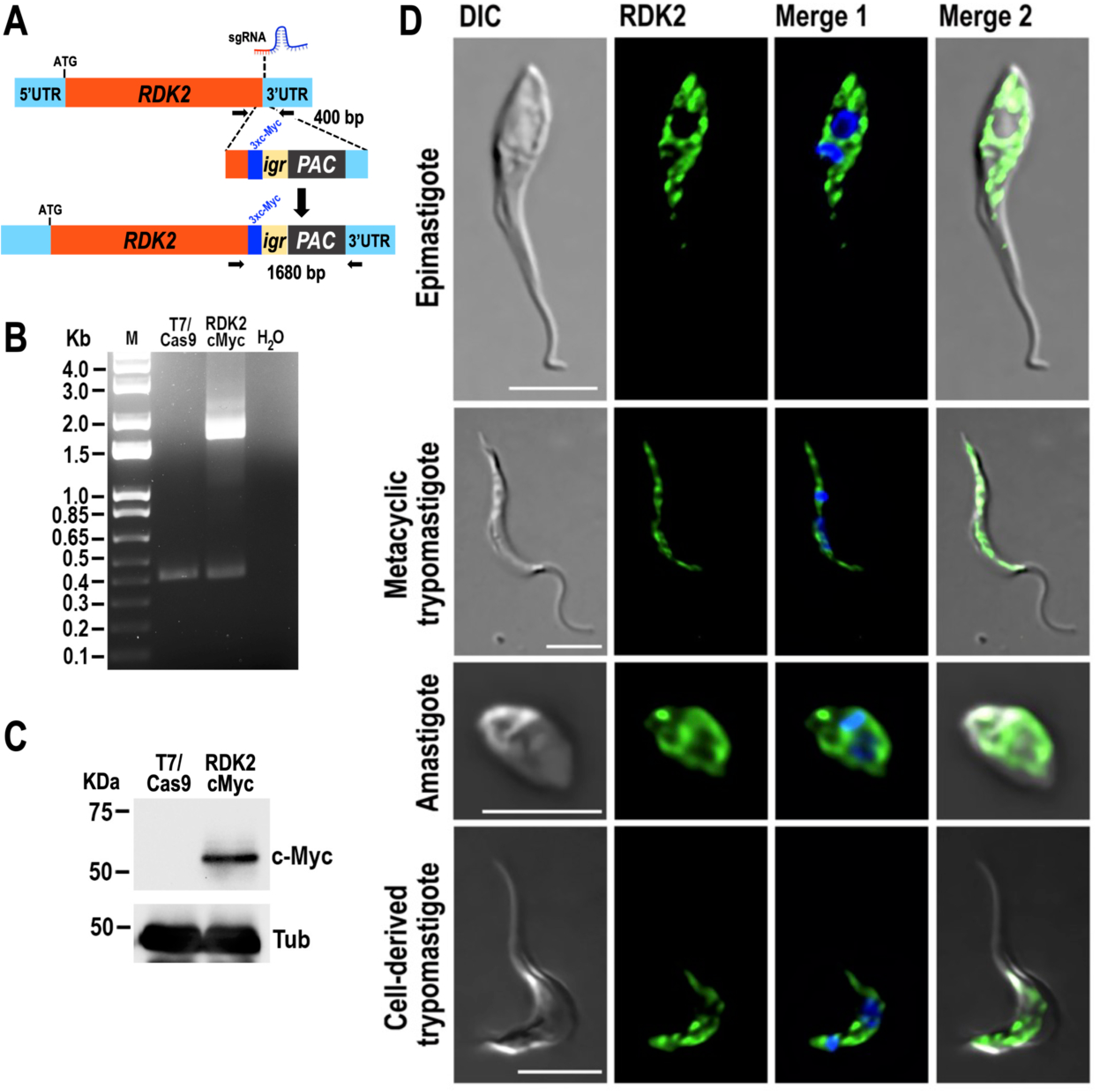
Analysis of expression of endogenously tagged *TcRDK2*-3xc-Myc. (A) Schematic representation of CRISPR/Cas9 mediated strategy for the C-terminal tagging of *TcRDK2.* The Homologous-directed repair determines integration of 3xc-Myc and antibiotic resistance gene (*Pac*, Puromycin N-acetyltransferase) at 3’ end of *TcRDK2.* UTR, untranslated region; *Igr*, tubulin intergenic region. Horizontal *black arrows* indicate primers used for tagging verification. (B) PCR verification of integration of DNA donor in the 3’ end of the endogenous *TcRDK2* locus*. TcRDK2* allele size: WT, 400 bp; 3xc-Myc tagged, 1680 bp. Lanes: M, 1-kb plus ladder; T7/Cas9, parental control; RDK2 cMyc, *TcRDK2*-3xc-Myc; H_2_O, PCR negative control. (C) *TcRDK2* endogenous tagging was verified by western blotting using antibodies anti-c-Myc (c-Myc). Tubulin (Tub) was used as a loading control. T7/Cas9, parental control, RDK2 cMyc, TcRDK2-3xc-Myc. (E) Immunofluorescence analysis (IFA) showed localization of endogenously tagged *TcRDK2*-3xc-Myc detected with anti-c-Myc antibodies (*green*) in epimastigote, Metacyclic trypomastigote, Amastigote, and Cell-derived trypomastigote. Merge of *green* signal (RDK2) and DAPI staining (*blue*), and DIC images are also shown. Scale bars = 5 μm.

### *TcRDK2* ablation affects metacyclogenesis, nucleus/kinetoplast configuration and *in vitro* infection of host cells

To explore the role of TcRDK2, we generated a *TcRDK2* knockout cell line using a CRISPR/Cas9/T7RNAP-mediated method as previously described (30, 31). This method involves co-transfecting epimastigotes that constitutively express T7RNAP and Cas9 with a sgRNA template and two donor DNAs, to replace both alleles by homologous-directed repair (HDR) with two resistance markers (*PAC* and *BSD*) (Fig. 2A). Stable transfectants were confirmed by PCR in clonal populations (Fig. 2A and B). We then studied the effects of *TcRDK2* deletion on proliferation, differentiation and infectivity of *T. cruzi*. *TcRDK2*-KO epimastigotes exhibited no significant difference in growth rate relative to the parental cell line when cultured in LIT medium (Fig. 2C and S3A). However, the ability of *TcRDK2*-KO epimastigotes to differentiate *in vitro* into metacyclic trypomastigotes (metacyclogenesis) was significantly reduced compared to that of control cells (Fig. 2D). To further characterize the phenotypic defects associated with *TcRDK2* knockout, we quantified the number of nuclei and kinetoplasts per cell after DAPI staining in control and *TcRDK2*-KO epimastigotes grown in LIT medium (Fig. 2E). During the normal cell cycle of *T. cruzi* epimastigotes, the nuclear/kinetoplast (N/K) configuration progresses from 1N1K to 1N2K and then to 2N2K; the kinetoplast divides before the nucleus, and mitosis of 1N2K cells generates 2N2K cells, which subsequently undergo cytokinesis to yield two 1N1K cells (32). Relative to controls, *TcRDK2*-KO epimastigotes showed a higher proportion of binucleated cells, with a significant increase in 2N1K cells. We also observed a significant accumulation of cells with abnormal numbers of nuclei and kinetoplasts (XNXK) (Fig. 2E and F). Together, these observations are consistent with a defect in cytokinesis.

**Figure 2.**
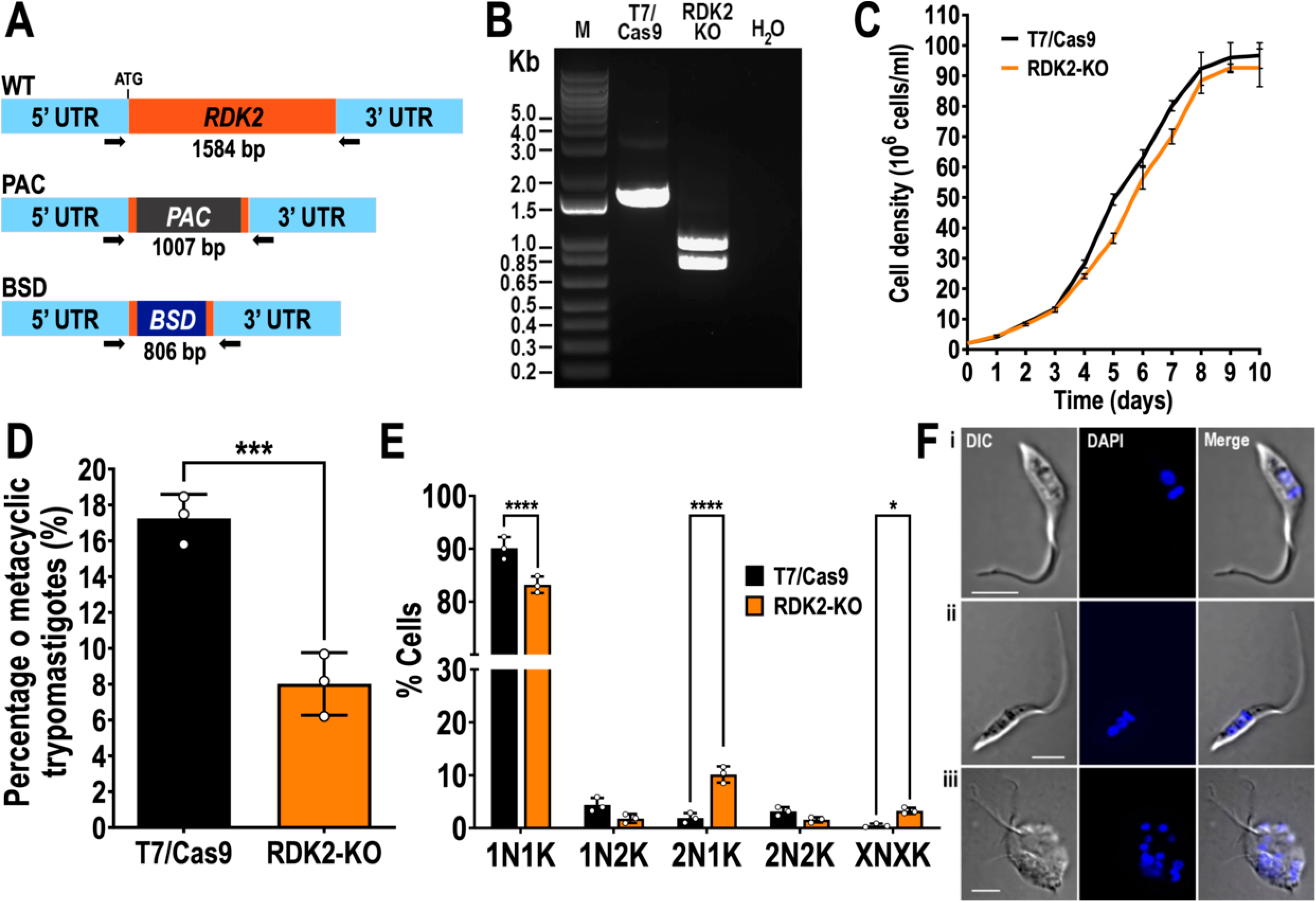
Generation and characterization of *TcRDK2*-KO cells. (A) Schematic representation of PCR primers used to verify *TcRDK2* ablation. Arrows indicate primers added to the reaction. PCR fragment sizes: WT *TcRDK2*, 1584 bp; *PAC*-replaced allele, 1007 bp, *BSD* (Blasticidin S deaminase)-replaced allele, 806 bp. (B) PCR verification of resistance cassette integration in both *TcRDK2* alleles. Lanes: M, 1-kb Plus DNA ladder; C, T7/Cas9, parental control; RDK2 KO, *TcRDK2*-KO; H_2_O, PCR negative control. (C) Growth of control (T7/Cas9) and *TcRDK2*-KO (RDK2-KO) epimastigotes in LIT medium (*n* = 3). (D) Percentage of metacyclic trypomastigotes in epimastigote cultures after incubation in TAU 3AAG medium. T7/Cas9 (control) and *TcRDK2*-KO (RDK2-KO) epimastigote differentiation to metacyclic trypomastigotes was quantified by staining with DAPI to distinguish the position of the kinetoplast by fluorescence microscopy. Values are means ± S.D. (*n* = 3), \*\*\**P* < 0.001 by Student’s *t*-test. (E) Effect of *TcRDK2* gene KO on nucleus/kinetoplast patterns established after DNA labeling with DAPI. At least 200 cells were counted for each analysis. Data represent means, and error bars indicate S.D. from *n* = 3 independent experiments. XNXK, X > 2. \**P* < 0.05; \*\*\*\**P* < 0.0001 (two-way ANOVA with Sidak’s multiple comparisons test). (F) Representative images of the main cell morphologies and DNA contents observed in *TcRDK2*-KO epimastigotes: (i) 1N1K, (ii) 2N1K, and (iii) XNXK. DIC, DAPI staining (blue) and merge images are shown. Scale bars = 5 μm.

We also assessed the capacity of *TcRDK2*-KO parasites to infect tissue-cultured cells. Repeated attempts to infect hFFs with purified metacyclic trypomastigotes were unsuccessful, as we did not detect egressed cell-derived trypomastigotes or intracellular amastigotes. In multiple independent infection experiments, starting around day 6 post-infection, we consistently observed a few host cells containing multiple vacuole-like structures with 1–4 *T. cruzi* parasites each, which did not resemble any defined developmental stage. These parasites remained motile within the vacuoles but failed to undergo normal egress or differentiate into trypomastigotes (Fig. S4). These findings indicate that *TcRDK2* is important for *T. cruzi* metacyclogenesis and is essential for infection of mammalian cells. Notably, attempts to generate add-back cell lines by overexpressing TcRDK2 in *TcRDK2*-KO epimastigotes were unsuccessful (see below).

### Overexpression of full-length TcRDK2 and a PH domain-truncated variant impairs growth and promotes the differentiation of *T. cruzi* epimastigotes

To evaluate the effect of TcRDK2 overexpression throughout the *T. cruzi* life cycle, we transfected epimastigotes with constructs harboring the *TcRDK2* ORF cloned into the pTREXn-3xHA and pTREXp-3xc-Myc expression vectors. However, attempts to obtain *TcRDK2*-overexpressing transfectants were unsuccessful, which may also explain why we were unable to generate *TcRDK2* add-back cell lines. To overcome this limitation, we used a tetracycline-inducible system (see *Methods*) to allow a controlled elevation of TcRDK2 levels and to minimize selection against clones with chronically high toxic kinase expression. This system enabled the analysis of acute effects of TcRDK2 overexpression at defined life-cycle stages. We used both the full-length kinase (RDK2^WT^) and a C-terminal PH domain-truncated variant (RDK2^ΔPH^) to assess the contribution of this domain to TcRDK2 function (Fig. 3A). Overexpression of RDK2^WT^ and RDK2^ΔPH^ epimastigotes was confirmed after induction with 0.5 μg/ml tetracycline by western blot (Fig. 3B) and IFA in epimastigotes (Fig. 3C). Interestingly, the subcellular localization of both RDK2^WT^ and RDK2^ΔPH^ was similar to that observed in the endogenously tagged *TcRDK2* cell lines (Fig. 1D and S2D), suggesting that neither overexpression nor loss of the PH domain appreciably alters the distribution of the protein within the cell.

**Figure 3.**
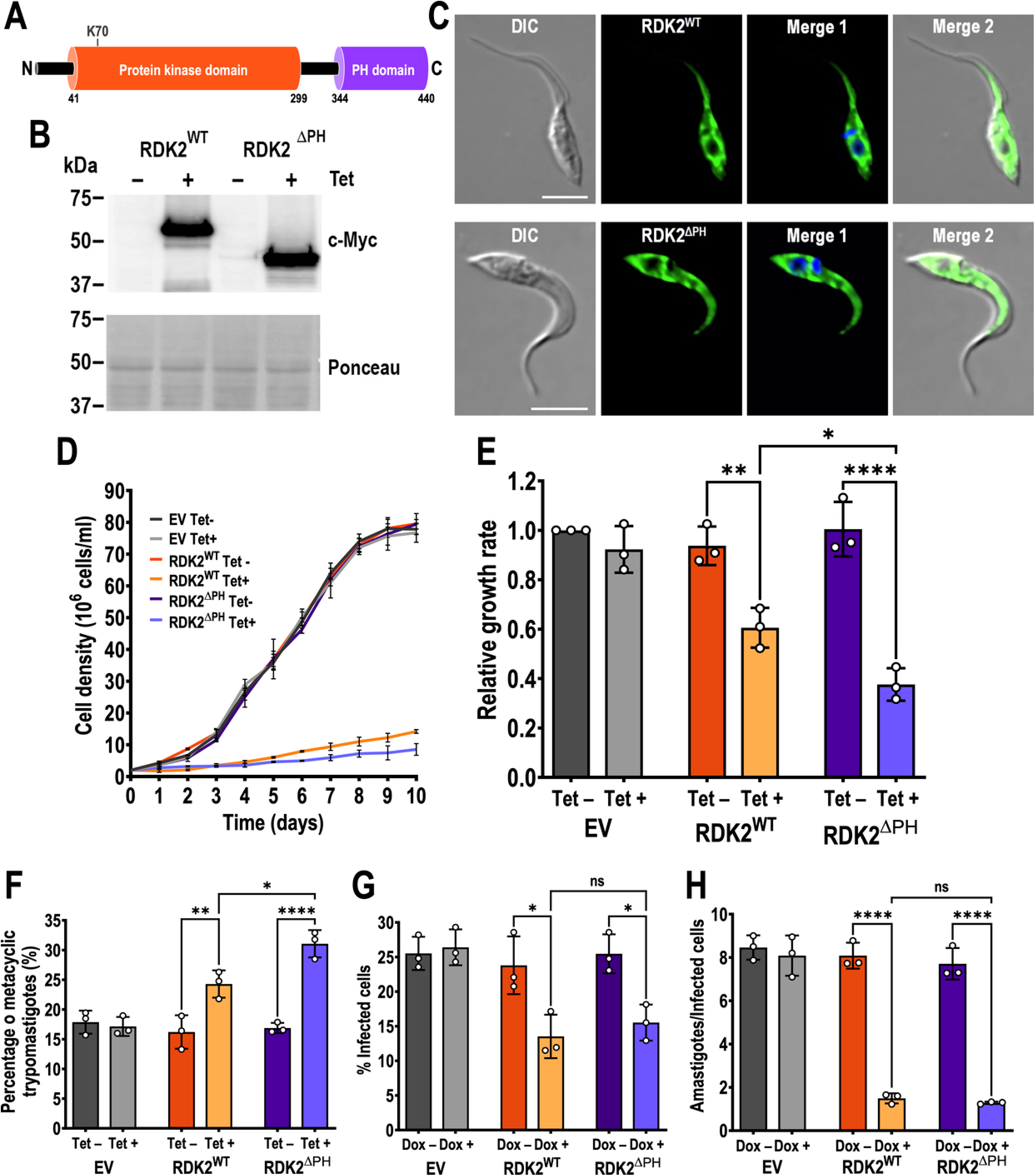
Phenotypic effects of overexpression of TcRDK2. *T. cruzi* epimastigotes were transfected with pTcINDEX-*RDK2*^WT^ (RDK2^WT^, full-length *TcRDK2*) and pTcINDEX-*RDK2*^ΔPH^ (RDK2^ΔPH^, truncated C-terminal PH domain). (A) Schematic representation of the TcRDK2 topology. The predicted amino acid sequence of *TcRDK2* (TriTrypDB ID: TcYC6_0103940) was scanned by InterProScan (https://www.ebi.ac.uk/interpro/) to search for protein functional domains. Serine/threonine protein kinase catalytic domain (InterPro IPR000719); Pleckstrin homology (PH) domain (InterPro IPR001849). Number under each domain indicate amino acids spanning these domains in TcRDK2. The position of the conserved lysine (K70) of the catalytic domain is shown. N- and C-terminus are indicated. (B) Western blot analysis of total protein extracts of epimastigotes transfected with RDK2^WT^ and RDK2^ΔPH^ in the absence (Tet−) or presence (Tet+) of 0.5 μg/ml tetracycline for 24 h using anti-c-Myc antibodies. Ponceau red staining is shown as loading control. (C) Immunofluorescence analysis (IFA) showed localization of RDK2^WT^ and RDK2^ΔPH^ in the presence of 0.5 μg/ml tetracycline for 24 h. Tagged RDK2^WT^-3xc-Myc (top panel) and RDK2^ΔPH^-3xc-Myc (bottom panel) were detected with anti-c-Myc antibodies (*green*) in epimastigote, Merge images are also shown. Scale bars = 5 μm. (D) Growth curve of EV (empty vector, control), RDK2^WT^ and RDK2^ΔPH^ epimastigotes in the absence (Tet−) or presence (Tet+) of 0.5 μg/ml tetracycline. Growth rate (E) was estimated between days 3 to 7 and were normalized using the EV Tet− condition as the control group. (F) Percentage of metacyclic trypomastigotes in EV (empty vector, control), RDK2^WT^ and RDK2^ΔPH^ epimastigote cultures after incubation in TAU 3AAG medium in the absence (Tet−) or presence (Tet+) of 0.5 μg/ml tetracycline. (G) EV (empty vector, control), RDK2^WT^ and RDK2^ΔPH^ trypomastigote infection of hFF cells. Cell-derived trypomastigotes were analyzed in the absence (Dox−) or pre-treated with 1.0 μg/ml doxycycline (Dox+) for 2 h before infection and during the invasion. The percentage of infected cells was assessed 4 hours post infection by fluorescence microscopy. (H) Intracellular replication of EV (empty vector, control), RDK2^WT^ and RDK2^ΔPH^ amastigotes analyzed in the absence (Dox−) or pre-treated with 1.0 μg/ml doxycycline (Dox+) for 48 h post infection. In panels D to H values are means ± S.D. (*n* = 3). For the results shown in panels E and H, statistical analysis was performed using two-way ANOVA with Tukey’s multiple comparisons test, whereas for panels F and G two-way ANOVA with Sidak’s multiple comparisons test was used. \**P* < 0.05; \*\**P* < 0.01; \*\*\*\**P* < 0.0001; ns, not significant. For the results shown in panels E–H, only biologically relevant pairwise comparisons are indicated.

To evaluate the effect of overexpression of RDK2^WT^ and RDK2^ΔPH^ on epimastigote replication, we performed growth curves of control (EV, *T. cruzi* epimastigotes transfected with pTcINDEX/3xTetO/3xc-Myc empty vector), RDK2^WT^, and RDK2^ΔPH^ epimastigotes in the absence (Tet−) or presence (Tet+) of tetracycline (Fig. 3D). Significant lower growth rates were observed for RDK2^WT^ and RDK2^ΔPH^ under tetracycline induction compared to the corresponding uninduced cultures and with EV cells in both conditions (Fig. 3D and E). No morphological alterations were detected in the induced mutants. This phenotype indicates that TcRDK2 overexpression impairs epimastigote replication and this could explain why we were unable to obtain viable constitutive TcRDK2 overexpression lines. Moreover, RDK2^ΔPH^ cells exhibited also a significant lower growth rate compared with RDK2^WT^ epimastigotes in the presence of tetracycline (Fig. 3E). We next evaluated the effect of RDK2^WT^ and RDK2^ΔPH^ overexpression in the ability of epimastigotes to differentiate into metacyclic trypomastigotes in TAU 3AAG medium. As shown in Figure 3F, induced RDK2^WT^ and RDK2^ΔPH^ parasites exhibited a significant higher percentage of metacyclic trypomastigotes compared with controls. Likewise, induction of RDK2^ΔPH^ overexpression (Tet+ condition) showed a significant higher percentage of metacyclic trypomastigotes compared to induced RDK2^WT^ cells. We also wanted to determine whether overexpression of RDK2^WT^ and RDK2^ΔPH^ in epimastigotes cultured in LIT medium affects differentiation into metacyclic trypomastigotes (Fig. S5A). After 120 h under Tet− or Tet+ conditions, tetracycline-induced RDK2^WT^ and RDK2^ΔPH^ populations displayed a significantly higher proportion of both intermediate forms (cells in which kinetoplast repositioning from the mid-cell region near the flagellum to the posterior end was incomplete) and fully differentiated metacyclic trypomastigotes than their respective controls. Consistent with the results in TAU 3AAG medium, induced RDK2^ΔPH^ parasites differentiated significantly more than induced RDK2^WT^ cells. Together, these findings indicate that *TcRDK2* positively modulates metacyclogenesis in *T. cruzi* epimastigotes.

### TcRDK2 overexpression compromises *T. cruzi* infection of mammalian cells

We also examined how overexpression of RDK2^WT^ and RDK2^ΔPH^ affects *T. cruzi* infection of host cells. Briefly, for the invasion assay, cell-derived trypomastigotes were pre-incubated with or without doxycycline for 2 h and maintained under the same conditions during the invasion period of hFF cells. After 4 h, trypomastigotes were washed out, and fresh medium without doxycycline was added to the cells, that were then incubated for 24 h before fixation and DAPI staining on coverslips for microscopy analysis. A significantly lower number of infected hFF cells was observed for both RDK2^WT^ and RDK2^ΔPH^ overexpression parasites under doxycycline induction compared to the uninduced parasites (Fig. 3G). We also evaluated the effect of TcRDK2 overexpression on the intracellular proliferation of amastigotes. For this, hFF cells were first infected with uninduced cell-derived trypomastigotes. After a 4h invasion period, extracellular parasites were removed, and cell cultures were maintained under Dox– or Dox+ conditions for 72 h. Then, cells were microscopically examined to determine the average number of intracellular amastigotes per infected cell. As observed for host-cell invasion, intracellular amastigote replication was also significantly reduced in both RDK2^WT^ and RDK2^ΔPH^ induced parasites (Fig. 3H). Overall, these experiments show that *TcRDK2* overexpression, whether full-length or ΔPH, markedly impairs host-cell invasion and subsequent intracellular amastigote proliferation.

### Overexpression of catalytically inactive RDK2 has a dominant-negative effect throughout *T. cruzi* life cycle

We set out to determine whether a catalytically inactive variant could also serve as a tool to elucidate the role of *TcRDK2* in the life cycle of *T. cruzi*. A common approach to generating catalytically inactive kinases is to mutate critical conserved residues within the kinase core. Indeed, substituting the conserved lysine of the catalytic domain has been repeatedly used to produce a “dead” kinase (33–35). The conserved Lys of a protein kinase catalytic domain is essential for kinase activity, by anchoring and orienting the α- and β-phosphates of ATP. This residue also stabilizes the active conformation by forming a salt bridge with a glutamic acid residue on the C-helix, which is crucial for the phosphotransfer reaction (36). For this purpose, we identified the conserved Lys residue in the catalytic domain of TcRDK2. As shown in the Alphafold 3.0-based structural model (Fig. 4A), Lys70 forms interactions with the ATP molecule and with Glu88 within the cleft-like pocket located between the small N-terminal lobe and the C-terminal lobe of TcRDK2. To this end, Lys70 was substituted by Ala to generate the TcRDK2^K70A^ variant, which was cloned into pTcINDEX/3xTetO/3xc-Myc and transfected into *T. cruzi* epimastigotes to establish an inducible TcRDK2^K70A^ overexpression cell line. Expression of TcRDK2^K70A^ upon tetracycline induction was confirmed by western blot analysis (Fig. 4B). Next, to assess the impact of RDK2^K70A^ overexpression on epimastigote proliferation, by monitoring the growth of control (EV) and RDK2^K70A^ parasites in the absence (Tet–) or presence (Tet+) of tetracycline (Fig. 4C). Although growth rates during the exponential phase of growth were similar (Fig. S3B), induced RDK2^K70A^ epimastigotes reached a significantly lower cell density at stationary phase compared to uninduced parasites and EV (Tet–/+) control epimastigotes (Fig. 4C). We also analyzed the nuclear/kinetoplast (N/K) configurations of RDK2^K70A^ epimastigotes in exponential phase, cultured in LIT medium with or without tetracycline, observing a significant increase in 2N1K cells (Fig. 4D). We next examined the effect of RDK2^K70A^ overexpression on *in vitro* metacyclogenesis in TAU 3AAG medium. Induced RDK2^K70A^ parasites showed a significant decrease in metacyclogenesis relative to controls (Fig. 4E), supporting a role for *TcRDK2* in metacyclogenesis of *T. cruzi* epimastigotes. However, epimastigotes overexpressing RDK2^K70A^ that were aged in LIT medium did not show defects compared with the controls (Fig. S5B). We also evaluated how RDK2^K70A^ overexpression affects *T. cruzi* during host-cell infection. Using the assay described above, both host-cell invasion and intracellular amastigote replication were significantly impaired in RDK2^K70A^ induced parasites compared to uninduced and with EV cells under either condition (Fig. 4F and G). Collectively, these results closely resembled those observed with the *TcRDK2*-KO cell line, suggesting that the K70A mutation abolishes RDK2 catalytic activity and thereby accounts for the observed defects.

**Figure 4.**
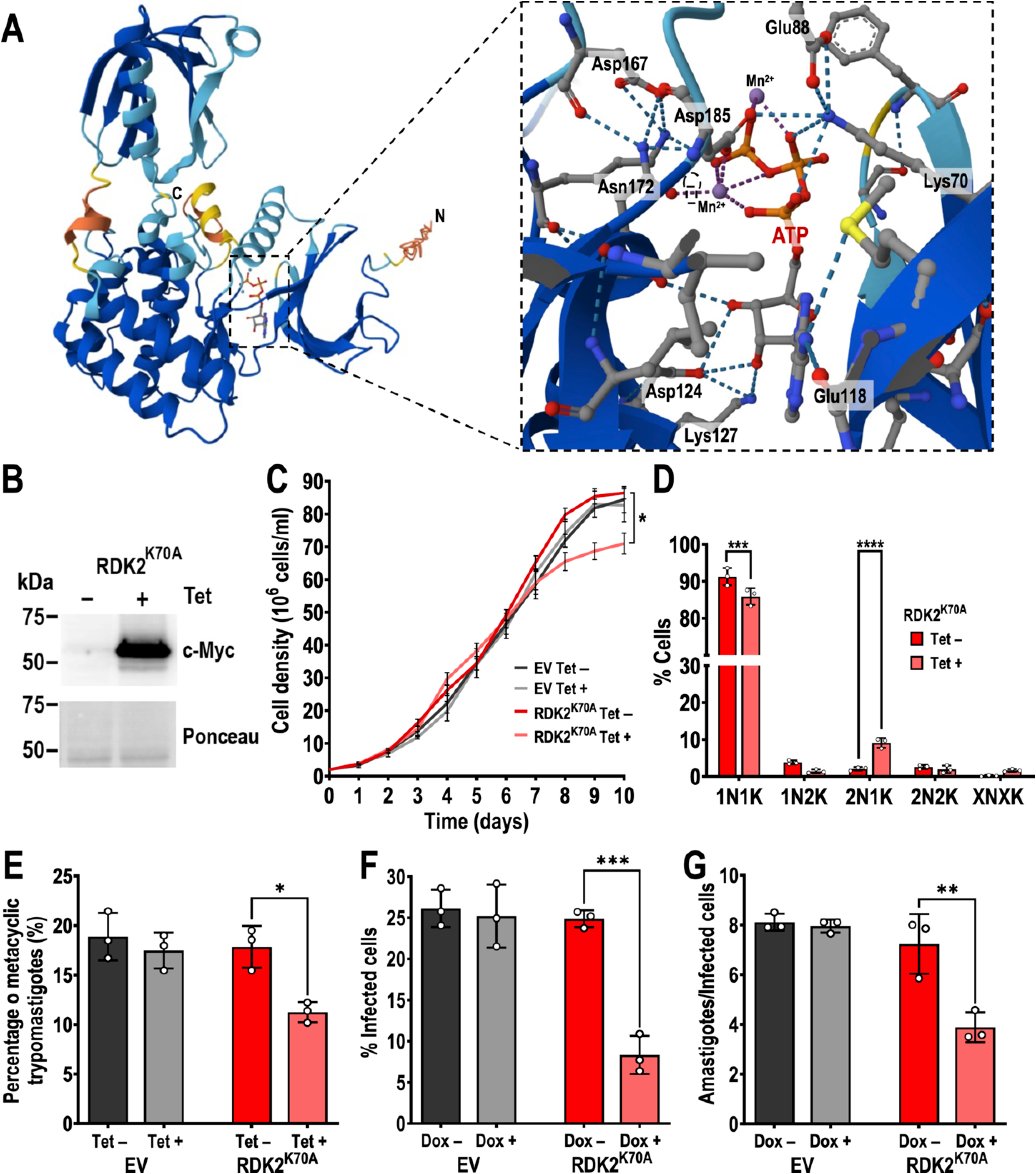
Phenotypic effects of overexpression of mutation of the conserved lysine in the catalytic domain of TcRDK2. *T. cruzi* epimastigotes were transfected with pTcINDEX-*RDK2*^K70A^, in which the predicted conserved lysine residue (K70) of the catalytic domain of TcRDK2 was substitute by alanine. (A) Cartoon representation of TcRDK2 with ATP binding in the catalytic domain made with AlphaFold 3. In a close-up is shown the kinase domain with interactions between ATP and crucial residues. Critical structures are highlighted in different colors and labeled by the sites. A divalent cation (Mn^2+^, purple balls) which is required for the catalysis reaction, was included for the modeling. In the close-up, it clearly shows that Glu88 forms a salt-bridge with Lys70 to hold the α- and β-phosphates in position. (B) Western blot analysis of total protein extracts of epimastigotes transfected with RDK2^K70A^ in the absence (Tet−) or presence (Tet+) of 0.5 μg/ml tetracycline for 24 h using anti-c-Myc antibodies. Ponceau red staining is shown as loading control. (C) Growth curve of EV (empty vector, control) and RDK2^K70A^ epimastigotes in the absence (Tet−) or presence (Tet+) of 0.5 μg/ml tetracycline. Two-way ANOVA with multiple comparisons was applied to compare the cell density from each growth curve at day 10. (D) Effect of RDK2^K70A^ overexpression on epimastigote nucleus/kinetoplast patterns in the absence (Tet−) or presence (Tet+) of 0.5 μg/ml tetracycline for 72 h. DNA was stained with DAPI. At least 200 cells were counted for each analysis. XNXK, X > 2. (E) Percentage of metacyclic trypomastigotes in EV (empty vector, control) and RDK2^K70A^ epimastigotes epimastigote cultures after incubation in TAU 3AAG medium in the absence (Tet−) or presence (Tet+) of 0.5 μg/ml tetracycline. (F) EV (empty vector, control) and RDK2^K70A^ trypomastigote infection of hFF cells. Cell-derived trypomastigotes were analyzed in the absence (Dox−) or pre-treated with 1.0 μg/ml doxycycline (Dox+) for 2 h before infection and during the invasion. The percentage of infected cells was assessed 4 hours post infection by fluorescence microscopy. (G) Intracellular replication of EV (empty vector, control) and RDK2^K70A^ amastigotes analyzed in the absence (Dox−) or pre-treated with 1.0 μg/ml doxycycline (Dox+) for 48 hours post infection. In panel C to G, data represent means, and error bars indicate S.D. from *n* = 3 independent experiments. For the results shown in panels C and G, statistical analysis was performed using two-way ANOVA with Tukey’s multiple comparisons test, whereas for panels D, E and F two-way ANOVA with Sidak’s multiple comparisons test was used. \**P* < 0.05; \*\**P* < 0.01; \*\*\**P* < 0.001; \*\*\*\**P* < 0.0001. For the results shown in panels E–G, only biologically relevant pairwise comparisons are indicated.

### Phosphoproteomic analysis of *TcRDK2*-overexpressing epimastigotes reveals the presence of potential substrates

To identify potential substrates of *T. cruzi* RDK2 and to characterize TcRDK2–induced changes in the cellular phosphoproteome, we conducted phosphoproteomic analysis on RDK2^WT^ epimastigotes after 48 h of tetracycline-induced overexpression. The experiment was performed including three biological replicates for each group: control (EV) and RDK2^WT^. Because one of the control samples generated an unrecoverable corrupted data file, the analysis was conducted using two control samples for comparison. Principal component analysis showed clear separation between the groups, supporting the use of this dataset for comparative analysis. The phosphoproteomic analysis identified a total of 3,770 phosphopeptides, of which 667 showed *P* < 0.05 (Table S3). Phosphopeptides were considered upregulated if they were detected in at least three RDK2^WT^ samples with a log_2_ fold change ≥ 1.5, whereas downregulated phosphopeptides were defined as those detected in at least two EV samples with a log_2_ fold change ≤ 1.5. Using these criteria, we identified 276 upregulated and 199 downregulated phosphopeptides, corresponding to 201 and 149 phosphoproteins, respectively (Table S4 and S5). Because many significantly dysregulated phosphorylation sites were detected in all replicates of only one condition (158 phosphopeptides corresponding to 142 proteins present exclusively in RDK2^WT^ samples and 134 phosphopeptides corresponding to 117 proteins present exclusively in EV samples (Table S6 and S7) conventional fold-change calculations were not suitable for comparing among each group. For phosphosites detected exclusively in RDK2^WT^ samples, we therefore used their normalized intensities within that group to calculate per-site Z-scores, visualized their relative abundance across biological replicates, and plotted the mean Z-score in a heat map (Fig. S6). Analogously, phosphosites detected only in EV samples were interpreted as RDK2^WT^-downregulated sites and analyzed in the same manner (Fig. S7). This set of upregulated phosphoproteins detected only in RDK2^WT^-induced replicates suggests that they may represent preferential substrates of TcRDK2, or components whose phosphorylation is altered as a downstream consequence of TcRDK2 activity. Interestingly, the top upregulated proteins included a small glutamine-rich tetratricopeptide repeat protein (SGT, TcYC6_0097790) and protein phosphatase 2C (PP2C, TcYC6_0018600), which were previously identified as key components whose expression and phosphorylation status, respectively, are modulated during *T. cruzi* metacyclogenesis (37). Moreover, several translation initiation factors were found to be significantly upregulated (eIF2B4, eIF2B5, eIF3c, eIF3E, eIF4E1, eIF4E3, eIF4G1, eIF4G2) and downregulated (eIF4G4, eIF2B, eIF4E3) phosphoproteins in our analysis, which suggests that TcRDK2 activity is linked to widespread remodeling of the translation initiation machinery. In addition, two basal body proteins were also found among the upregulated phosphoproteins, suggesting that TcRDK2 may influence kinetoplast segregation and cytokinesis at least in part by modulating phosphorylation of basal body-associated components. Importantly, ortholog searches in TriTrypDB revealed that the upregulated phosphoprotein set included 61 hypothetical proteins with no predicted function, whereas 20 hypothetical proteins were present in the downregulated group (Table S4 and S5). Seven phosphorylated residues, distributed across six phosphopeptides derived from the overexpressed TcRDK2, were also found to be upregulated in the phosphoproteomic analysis (Fig. 5A and B), although they were excluded from subsequent analyses. Among these residues, three (S189, S195, T207) are located within the predicted activation loop of TcRDK2, whereas the S341/S342 and S381 phosphoamino acids were identified immediately upstream of and within the PH domain, respectively. T207 may represent the key phosphorylated residue within the TcRDK2 activation loop, as it lies within the FCGT motif in this region. In NEK kinases, the conserved serine or threonine that serves as the principal activating site is typically positioned within an F/LxxS/T motif (27). We also identified several protein kinases (Table S8) among the dysregulated phosphoproteins, suggesting that TcRDK2 plays a role as a kinase network regulator.

**Figure 5.**
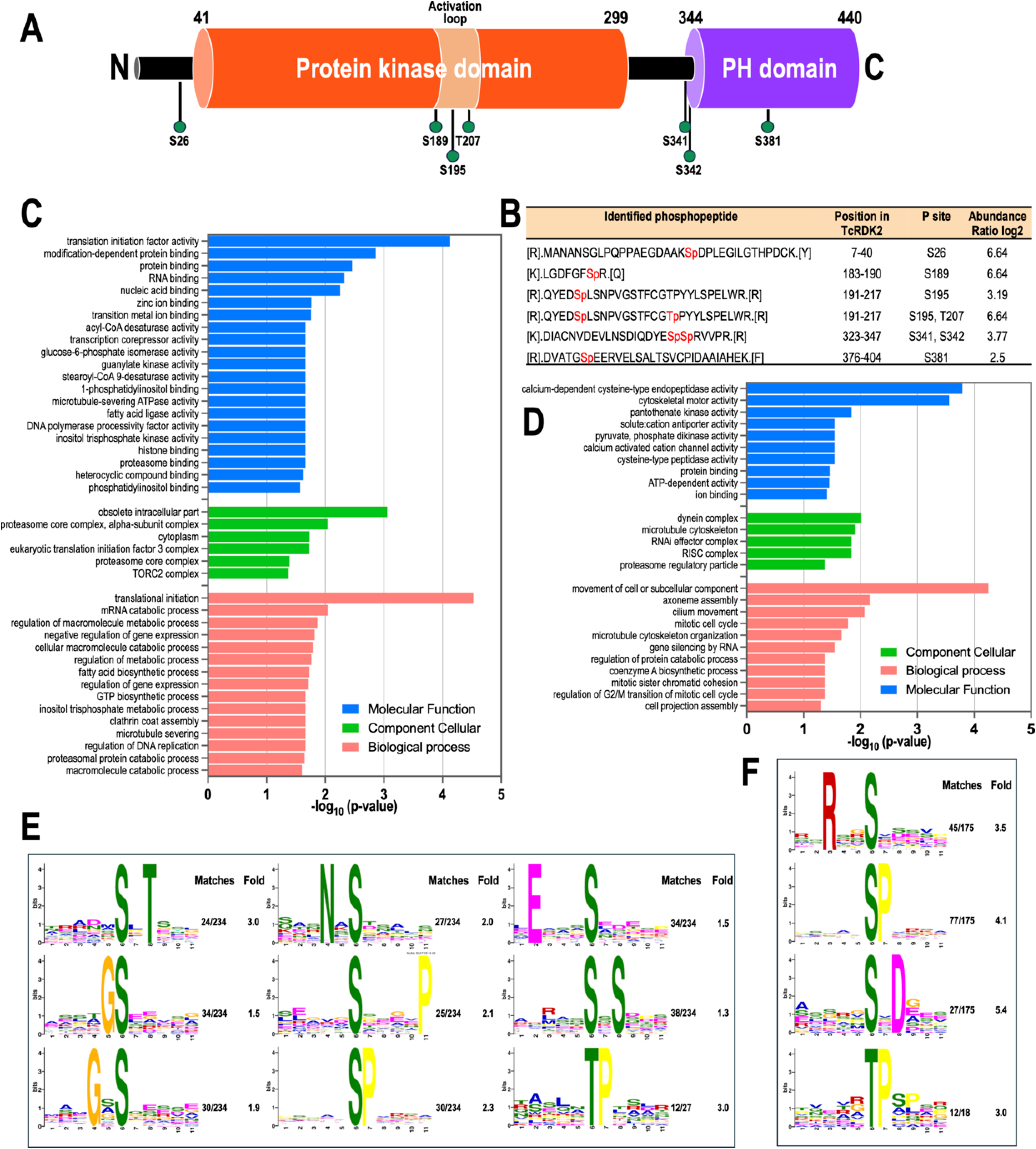
Phosphoproteomic analysis of *TcRDK2* overexpression. (A) Schematic representation of the TcRDK2 topology with highlighted significantly phosphorylated amino acid residue (green balls: S26, S189, S195, T207, S341, S342, and S381) that were identified by mass spectrometry in the RDK2^WT^ overexpression samples. Number above each domain indicate amino acids spanning these domains in TcRDK2. The predicted activation loop is highlighted (light orange) as part of the structure of the protein kinase domain. N- and C-terminus are indicated. (B) List of identified phosphopeptides of TcRDK2 after inducing RDK2^WT^ overexpression with tetracycline. In red (first column) are highlighted the phosphorylated S/T in each phosphopeptide. The abundance of phosphopeptides was determined by comparison with protein extracts from control epimastigotes (pTcINDEX empty vector). Phosphopeptides with a log_2_ fold change of 6.4 were detected exclusively in the RDK2^WT^ overexpression samples. (C and D) Gene ontology (GO) enrichment analysis of significantly upregulated (C) and downregulated (D) differentially expressed phosphopeptides (DEP) in RDK2^WT^ with predicted GO terms (*p* < 0.05). (E and F) Phosphorylation site motif analysis of the upregulated (E) and downregulated (F) DEPs using Motif-X. The degree of amino acids emerging at specific locations is represented by the elevation of the symbols. Matches represent the occurrences of phosphopeptides within a group. Fold indicates the relative enrichment of overrepresented sequence motifs as compared to the shuffled foreground peptides as background peptides.

A Gene Ontology (GO) term enrichment analysis was performed of upregulated and downregulated phosphoproteins in Tet-induced RDK2^WT^ parasites (Fig. 5C and D). For the upregulated phosphoproteins (Fig. 5C), translation initiation factor activity (GO:0003743), modification-dependent protein binding (GO:0140030), protein binding (GO:0005515), RNA binding (GO:0003723), and nucleic acid binding (GO:0003676) were the most abundant molecular function GO terms. Among the cellular component terms, obsolete intracellular part (GO:0005622), proteasome core complex (alpha-subunit complex) (GO:0005839), cytoplasm (GO:0005737), and eukaryotic translation initiation factor 3 complex (GO:0005852) were affected. The top four biological process GO terms found with the upregulated phosphoproteins were translational initiation (GO:0006413), mRNA catabolic process (GO:0006402), regulation of macromolecule metabolic process (GO:0060255), and negative regulation of gene expression (GO:0010629). We also sought to understand the potential functions of phosphoproteins that were downregulated upon RDK2^WT^ overexpression (Fig. 5D). Gene Ontology analysis indicated enrichment of several molecular function terms, including calcium-dependent cysteine-type endopeptidase activity (GO:0004198), cytoskeletal motor activity (GO:0003774), and pantothenate kinase activity (GO:0004594). In the cellular component category, the most prominent terms were the dynein complex (GO:0030286), microtubule cytoskeleton (GO:0015630), RNAi effector complex (GO:0031332), and RISC complex (GO:0016442). Among biological processes, the top enriched terms were movement of cell or subcellular component (GO:0006928), axoneme assembly (GO:0035082), cilium movement (GO:0003341), and the mitotic cell cycle (GO:0000278).

To investigate the specificity of TcRDK2-dependent phosphorylation, we used the MoMo webtool in combination with the motif-X algorithm to identify consensus substrate motifs within the −5 to +5 residues flanking the phosphosites of significantly dysregulated phosphopeptides. Of the phosphorylation sites included in this analysis, 261 were upregulated: 234 (89.7%) on serine and 27 (10.3%) on threonine. In contrast, 193 sites were downregulated, comprising 174 (90.7%) serine and 18 (9.3%) threonine residues. Nine motifs were identified among the upregulated phosphopeptides, eight of which were serine-centered (SxT, GS, GxS, NxS, SxxxxP, SP, ExxxS, SxS) and one threonine motif (TP) (Fig. 5E). Motif analysis of downregulated phosphopeptides in tetracycline-induced RDK2^WT^ cells compared with controls yielded three serine motifs (RxxS, SP, SxD) and one threonine motif (TP) (Fig. 5F).

## DISCUSSION

In this work we functional characterized the NIMA-related kinase (NEK) *TcRDK2* in *T. cruzi* and demonstrate that this kinase contributes to the regulation of epimastigote proliferation, kinetoplast division, metacyclogenesis, and infection of mammalian cells. Most importantly, our phosphoproteomic data indicates that overexpression of TcRDK2 cause a massive remodeling of the translation initiation machinery in *T. cruzi* epimastigotes.

NEK kinases are an evolutionarily conserved family of serine/threonine kinases originally identified in *Aspergillus nidulans* as NIMA (never in mitosis A), a key regulator of entry into mitosis (38). In opisthokonts, NEKs have diversified to perform multiple roles in cell-cycle progression, centrosome and spindle function, ciliogenesis, and regulation of microtubule dynamics (26–28). In trypanosomatids, NEKs have been linked to the regulation of morphogenesis, basal body duplication, kinetoplast segregation, differentiation and cytokinesis (14, 16, 29, 39), and comparative kinome analyses indicate that these parasites possess an expanded and divergent NEK repertoire relative to other eukaryotes (15).

Endogenous tagging at both the N- and C-termini showed that *TcRDK2* is expressed in all four major *in vitro* stages (epimastigote, metacyclic trypomastigote, amastigote, and cell-derived trypomastigote) and exhibits a punctate cytoplasmic distribution. A similar localization pattern has been reported for the *T. brucei* and *L. mexicana* RDK2 homologs (29, 40). The constitutive expression of this protein across developmental stages indicates that TcRDK2 is likely involved in core cellular processes required throughout *T. cruzi* life cycle.

Similar to canonical eukaryotic protein kinases, TcRDK2 contains an activation loop within its catalytic core. Notably, the predicted activation-loop threonine (T207) was found phosphorylated *in vivo*, consistent with the usual role of activation-loop phosphorylation in stabilizing the active conformation (41), a function that would be interesting to confirm experimentally.

In this study we generated a catalytic-dead TcRDK2 variant (TcRDK2^K70A^) by substituting the conserved lysine in the catalytic domain. Overexpression of TcRDK2^K70A^ reproduced most of the phenotypes observed in the *TcRDK2*-KO line: normal growth, an increased proportion of abnormal 2N1K cells, impaired metacyclogenesis, and marked defects in host-cell invasion and intracellular amastigote proliferation. These extensive similarities indicate that TcRDK2^K70A^ acts as a dominant-negative mutant that interferes with endogenous TcRDK2 function, therefore supporting the requirement for TcRDK2 kinase activity *in vivo*. Analogous dominant-negative mutants generated by substituting the conserved catalytic lysine have been well characterized for several other protein kinases, including *aPKCι*, *PKA*, and *ULK1* (33, 42, 43). Similarly, overexpression of a kinase-dead *TbNRKC* variant, a NEK-family kinase, in *T. brucei* procyclic cells produced phenotypes similar to those observed after RNAi-mediated knockdown (39). Loss of *TcRDK2* function, whether by gene knockout or expression of the *TcRDK2*^K70A^ mutant, resulted in a significant increase in 2N1K and other abnormal XNXK cells, indicative of impaired coordination between mitosis, kinetoplast segregation, and cytokinesis. However, in contrast to *T. brucei*, where RNAi-mediated downregulation of *TbRDK2* in bloodstream forms affects kinetoplast division, cytokinesis, and overall cell growth (14), *TcRDK2*-KO epimastigotes grew at rates comparable to control parasites in LIT medium. Moreover, in *L. mexicana*, *LmxRDK2* proved refractory to gene deletion in promastigotes, indicating that this kinase is likely required for parasite replication at least in that developmental stage (29). In *T. cruzi* epimastigote cultures in exponential phase of growth, most cells (>80%) display a 1N1K configuration under normal conditions, whereas a smaller subset (∼2-3%) shows 1N2K, indicating that kinetoplast division has occurred and the cells are in late G_2_ or early mitosis; the remaining cells (3%) present a 2N2K configuration, corresponding to parasites that have completed mitosis and are undergoing, or are about to undergo, cytokinesis (32). Defects in kinetoplast division and cytokinesis, often manifesting as a 2N1K phenotype, generally reflect a stalled or dysregulated cell cycle in which the nuclear and kinetoplast cycles become uncoupled. Accumulation of 2N1K and multinucleated cells without an immediate collapse of proliferation over the first 24-48 h has been observed after manipulating the expression of cell cycle kinases in *T. brucei* (*TbPLK, TbNRKC*) and of the High Mobility Group B protein (*TcHMGB*) in *T. cruzi*; however, in all these cases, the sustained cytokinesis defect ultimately causes a profound loss of proliferative capacity (39, 44–46). A possible explanation for why in *TcRDK2* loss-of-function conditions these cytological defects appear not to impact bulk population of *TcRDK2*-KO cells growth in rich LIT medium is that the 2N1K state could represents a delay rather than an irreversible block, allowing some cells managed to divide their kinetoplast or to complete an asymmetric division in which at least one viable daughter is produced despite abnormal kinetoplast segregation (47, 48).

Interestingly, we also identified two basal body–associated proteins among the upregulated phosphoproteins in *TcRDK2*-overexpressing parasites. In *T. brucei*, kinases that target the basal body are known to be crucial for correct kinetoplast segregation, and their downregulation leads to defects in kinetoplast division and cytokinesis (23, 44, 49). TcRDK2-dependent phosphorylation of basal body components therefore offers a plausible link between this kinase and the abnormal N/K configurations observed in *TcRDK2* loss-of-function mutants, and it will be important in future work to directly examine basal body duplication and segregation in these lines.

Deletion of TcRDK2 and overexpression of the kinase-dead RDK2^K70A^ variant both caused a significant reduction in *in vitro* metacyclogenesis, whereas inducible overexpression of either full-length TcRDK2 or the PH-domain-truncated variant significantly enhanced differentiation into metacyclic trypomastigotes. These reciprocal phenotypes strongly suggest that *TcRDK2* acts as a positive regulator of metacyclogenesis in *T. cruzi*. Interestingly, in a quantitative proteome and phosphoproteome analysis performed during *T. cruzi* metacyclogenesis (Dm28c strain), TcRDK2 was identified among the proteins whose abundance increase during differentiation (37). In the same study, two TcRDK2 phosphosites, S403 and S404, were found among the stronger upregulated sites. These residues correspond to S341 and S342 in the TcYC6 *TcRDK2* sequence and were likewise found upregulated in our phosphoproteomic analysis. Together, these observations support a role for *TcRDK2* in the metacyclogenesis program of *T. cruzi*. This function is particularly intriguing when contrasted with *T. brucei* RDK2, which behaves as a repressor of bloodstream-to-procyclic differentiation (14). Additionally, RDK2 is strongly upregulated in *T. brucei* metacyclic cells, as quantitative proteomics showed it to be the most up-regulated kinase at this stage (50). In both parasites, metacyclic trypomastigotes are non-dividing forms specialized for transmission to the mammalian host, but *T. brucei* metacyclic trypomastigotes arise in the salivary glands of the tsetse fly whereas *T. cruzi* metacyclic trypomastigotes develop in the hindgut of kissing bugs (51, 52). Moreover, *T. cruzi* metacyclic trypomastigotes are aggressive intracellular invaders, while *T. brucei* metacyclics remain extracellular in the bloodstream, protected by a dense VSG coat (52, 53). Therefore, the biological context in which RDK2 operates at the metacyclic stage is distinct in the two species. Finally, assessing metacyclogenesis *in vivo* in infected kissing bugs would be highly valuable to validate our *in vitro* findings in a physiological context.

*TcRDK2*-KO parasites were unable to establish productive infections in human fibroblasts. Metacyclic trypomastigotes derived from the knockout failed to generate typical intracellular amastigotes or egressed trypomastigotes; instead, only host cells contained vacuole-like structures harboring a few atypical, rare motile parasites that did not progress through the normal intracellular cycle. These observations indicate that TcRDK2 is required for one or more steps in the transition from metacyclic trypomastigote to replicative amastigote and subsequent egress. It must also be considered that the lack of infectivity may reflect incomplete or defective metacyclogenesis in the knockout, leading to metacyclics that are morphologically present but functionally incompetent. In line with this, parasites overexpressing the catalytic-dead RDK2^K70A^ mutant were still able to invade host cells but showed severely impaired intracellular proliferation, suggesting that in the intracellular environment the dominant-negative effect of RDK2^K70A^ is less detrimental than complete loss of *TcRDK2*, yet still sufficient to compromise the full infection cycle and underscoring the requirement of TcRDK2 kinase activity for successful infection.

Inducible overexpression of TcRDK2 revealed that kinase dosage must be tightly controlled. Overexpression of either RDK2^WT^ or the PH-truncated RDK2^ΔPH^ dramatically slowed epimastigote proliferation, enhanced metacyclogenesis, and severely impaired host-cell invasion and intracellular amastigote proliferation, even though no obvious mislocalization was detected by IFA. The consistently stronger phenotypes of RDK2^ΔPH^ suggest that the PH domain normally restrains TcRDK2, either by limiting access to substrates, restricting subcellular distribution, or maintaining the kinase in a partially autoinhibited state, such that its removal unleashes more promiscuous or mis-timed phosphorylation (54–57).

A distinctive feature of TcRDK2 is its C-terminal PH domain, a modular element shared with several trypanosomatid NEKs but absent from NEKs in other organisms, suggesting that this kinase family has acquired lineage-specific regulatory modules that could be exploited for selective therapeutic targeting (15). PH domains typically engage phosphoinositides and/or protein partners at membranes and often regulate kinase localization or activity (58, 59). In *T. cruzi*, PH-like domains have also been predicted or described in other signaling kinases, including AGC family members (60, 61), where they are thought to mediate membrane recruitment and responsiveness to lipid signaling.

In metazoan AGC kinases such as Akt/PKB, the PH domain binds membrane phosphoinositides (e.g., PI(3,4,5)P₃ and PI(3,4)P₂), disrupting an intramolecular interface that keeps the kinase domain autoinhibited and thereby promoting activation-loop phosphorylation and full catalytic activation (57, 62). *T. cruzi* possesses the enzymatic machinery to synthesize PI(3,4,5)P₃, PI(3,4)P₂, PI(3,5)P₂, and PI(4,5)P₂ (63), and recent work on a *T. cruzi* Akt-like kinase has begun to highlight PIP-dependent regulation of protein kinases in this parasite (61). By analogy, the PH domain of TcRDK2 may integrate lipid-signaling inputs to control when and where this NEK is activated. The *T. cruzi* YC6 genome encodes ∼9 NEK kinases with predicted C-terminal PH domains (tritrypdb.org), yet the regulatory roles of these domains in NEK function have not been investigated. Dissecting whether TcRDK2 and other PH-containing NEKs respond to specific phosphoinositides, acting as lipid-regulated switches for cell-cycle progression and differentiation, will be an important next step toward understanding how TcRDK2 activity is controlled.

Phosphoproteomic profiling of *TcRDK2*-overexpressing epimastigotes revealed extensive remodeling of the phosphoproteome, with hundreds of up- and downregulated phosphosites. Among the stronger upregulated sites were those on SGT and PP2C, both previously shown to undergo expression or phosphorylation changes during metacyclogenesis (37), reinforcing a connection between TcRDK2 and differentiation-associated signaling. The prominent regulation of translation initiation factors (eIF2B, eIF3, eIF4 family members) suggests that TcRDK2 influences protein synthesis, consistent with observations that NEKs and other cell-cycle kinases can modulate translation in response to environmental or cell-cycle cues (26, 27). In addition, the GO term enrichment analyses suggest that TcRDK2 influences both the post-transcriptional control of gene expression and the organization of cytoskeletal and motility structures that are central to *T. cruzi* growth, morphogenesis, and infectivity. The presence of several kinase and phosphatase hits among TcRDK2-dependent phosphoproteins further suggests that TcRDK2 occupies an upstream or central position in a network of kinase-kinase and kinase-phosphatase interactions.

Motif analysis identified multiple serine- and threonine-centered motifs but no single dominant TcRDK2 consensus, implying that TcRDK2 acts within a broader phosphorylation network involving other kinases and phosphatases. Some motifs (e.g., SP) are associated with proline-directed kinases (64), while others (e.g., ExxxS) may be compatible with acidic/casein-like kinases (65). Although comprehensive substrate profiling indicates that many NEK kinases prefer hydrophobic residues at position -3 (e.g., FxxS/T or F/LxxS/T), it is likely that TcRDK2, like other NEKs, displays relatively broad substrate specificity with only modest amino-acid sequence constraints around the phosphorylation site (27, 66). Dissecting which of these motifs represent direct TcRDK2 targets will require *in vitro* kinase assays, *in vivo* analog-sensitive approaches, and systematic phosphosite mutagenesis. Because we did not acquire a matching global proteome, we cannot fully exclude that some phosphopeptide changes reflect altered protein abundance; nevertheless, the coordinated regulation of functionally related phosphoproteins (e.g., multiple eIFs) strongly supports a TcRDK2-dependent remodeling of phosphorylation networks. Moreover, after 48 h of *TcRDK2* overexpression, it is likely that a substantial fraction of the observed phosphorylation changes is mediated indirectly through downstream kinases. It will therefore be important to perform shorter time-course experiments to capture earlier, more direct TcRDK2-dependent events.

Finally, the large proportion of hypothetical proteins in *T. cruzi* is a well-recognized feature of its genome (67), and the substantial number of such proteins showing TcRDK2-dependent phosphorylation further highlights how much of the *T. cruzi* signaling landscape remains uncharacterized.

Taken together, our data position *TcRDK2* as a NEK-family kinase that regulates cell-cycle progression, kinetoplast segregation, metacyclogenesis, and host-cell infection in *T. cruzi*. The combination of (i) conservation across kinetoplastids, (ii) lineage-specific domain architecture, (iii) marked sensitivity to both loss and overexpression, and (iv) essentiality for infection and intracellular replication in mammalian cells poses *TcRDK2* as an attractive, *T. cruzi*-specific, potential drug target for Chagas disease treatment. Further work using analog-sensitive *TcRDK2* alleles or inducible gene-downregulation systems (to more precisely evaluate *TcRDK2* loss of function in host cells), together with time-resolved phosphoproteomics and analysis of gene expression profiles, will be essential to define the immediate substrates and signaling pathways controlled by TcRDK2.

## MATERIAL AND METHODS

### Chemicals and reagents

Fetal bovine serum (FBS) was purchased from R&D Systems (Minneapolis, MN). G418 was obtained from KSE Scientific (Durham, NC). Puromycin, blasticidin S HCl, Hygromycin B, Subcloning Efficiency DH5a competent cells, Pierce ECL Western blotting substrate and BCA Protein Assay Kit, Pierce Phosphatase inhibitor, Horseradish peroxidase (HRP)-conjugated anti-mouse IgG antibodies, mouse anti-c-Myc monoclonal antibody (9E10), and mouse anti-HA monoclonal antibody were from Thermo Fisher Scientific (Waltham, MA). Alexa Fluor 488-conjugated donkey anti-mouse conjugated donkey anti-rabbit was from Jackson ImmunoResearch (West Grove, PA). Restriction enzymes, NEBuilder^®^ HiFi DNA Assemble Master Mix and Q5^®^ High-Fidelity DNA Polymerase were obtained from New England BioLabs (Ipsich, MA). ZymoPURE Plasmid Miniprep, ZymoPURE II Plasmid Midiprep and DNA Clean & Concentrator-5 were from Zymo Research (Irvine, CA). T4 DNA Ligase and GoTaq G2 Flexi DNA Polymerase were from Promega (Madison, WI). cOmplete™ Mini EDTA-free Protease Inhibitor Cocktail was from Roche (Basel, Switzerland). 4-mm electroporation cuvettes, Precision Plus Protein Dual Color Standards, polyacrylamide and nitrocellulose membranes were from Bio-Rad (Hercules, CA). Fluoromount-G mounting medium was from Southern Biotech (Birmingham, AL). The pMOTag23M vector (68) was a gift from Dr. Thomas Seebeck (University of Bern, Bern, Switzerland). DNA oligonucleotides were purchased from Integrated DNA Technologies (Coralville, IA). Phenylmethylsulfonyl fluoride (PMSF), N-*p*-tosyl-L-phenylalanine chloromethyl ketone (TPCK), trans-epoxysuccinyl-l-leucylamido-(4-guanidino) butane (E64), protease inhibitor cocktail for use with mammalian cell and tissue extracts (Cat. No. P8340), Benzonase nuclease, and all other reagents of analytical grade were from Sigma-Aldrich (St. Louis, MO).

### Cell culture

*T. cruzi* Y strain epimastigotes were cultured in liver infusion tryptose (LIT) medium supplemented with 10% heat-inactivated fetal bovine serum (FBS) at 28°C (69), supplemented with 10% heat-inactivated fetal bovine serum (FBS), penicillin (100 I.U./mL), and streptomycin (100 µg/mL) at 28°C. According to their antibiotic resistance, mutant cell lines were maintained in medium containing, alone or in combination, 250 μg/ml G418, 5 μg/ml puromycin, 10 μg/ml blasticidin and 150 μg/ml Hygromycin. T7/Cas9 and T7/TetR cell lines were maintained in medium containing 250 μg/ml G418. Cell density was determined using a Guava^®^ Muse^®^ Cell A10 analyzer (Luminex Corporation, Austin, TX). Tissue culture-derived trypomastigotes and amastigotes were collected from the culture medium of infected human foreskin fibroblasts (hFFs) cells. hFFs were grown in DMEM (Dulbecco’s Modified Eagle Medium, Gibco) supplemented with 10% FBS, penicillin (100 I.U./mL), and streptomycin (100 µg/mL), and maintained with 5% CO_2_ at 37°C.

### In silico analysis

Nucleotide sequences of *T. cruzi* Y strain *TcRDK2* (TcYC6_0103940) gene was retrieved from TriTrypDB genome data base (tritrypdb.org). Prediction of protein domains of TcRDK2 amino acid sequences was made using web servers: https://ebi.ac.uk/interpro/ and https://prosite.expasy.org/scanprosite/. Selection of protospacers for knockout and tagging strategies was performed using EuPaGDT (eukaryotic pathogen CRISPR guide RNA/DNA design tool; http://grna.ctegd.uga.edu) (70). Prediction and modeling of TcRDK2 was performed by AlphaFold 3 (alphafoldserver.com).

### *TcRDK2* overexpression

*TcRDK2* open reading frame (ORF) (1,320 nt) was PCR amplified using Q5^®^ High-Fidelity DNA polymerase and *T. cruzi* Y strain genomic DNA as template (primers 1 and 2; Table S1), then it was cloned into the pTREXn/3xHA and pTREXp/3xc-Myc vector (71) by restriction sites XbaI/EcoRV. Gene cloning was confirmed by restriction analysis and DNA sequencing, and constructs were subsequently used to transfect *T. cruzi* epimastigotes.

### Generation of an inducible expression system for *T. cruzi*

For the inducible expression of *TcRDK2* we first generated a *T. cruzi* cell line constitutively expressing T7 RNA polymerase and the Tet repressor (TetR) after transfecting with the pTREXn/T7-TetR plasmid. Briefly, the coding sequence of the T7 RNAP and the TetR were amplified (primers 3 to 8; Table S1) using Q5^®^ High-Fidelity DNA polymerase from the pLEW13 plasmid, while the *T. cruzi* HX1 region was amplified from the pTREXn (72). Subsequently the three fragments inserted into pTREXn linearized with XbaI/HindIII using the NEBuilder^®^ HiFi DNA Assemble Master Mix downstream the rDNA promoter and a first copy of the HX1 region in the following order: T7RNAP-HX1-TetR. Finally, the pTREXn/T7-TetR plasmid was transfected in *T. cruzi* Y strain epimastigotes, and after selection with G418, a resistant clone was obtained by limiting dilutions, which was named as T7/TetR cell line. Later we generated a modified version of the tetracycline-regulated expression vector for *T. cruzi* pTcINDEX (73), in which we incorporated two additional Tet operator (TetO) regions, additional restriction sites in the multiple cloning sites (MCS) region and 3 copies of the c-Myc tag. Briefly, Q5^®^ High-Fidelity DNA polymerase was used to amplify 2xTetO, MCS and 3xc-Myc fragments using pLEW100 vector, pTcINDEX-c-Myc and pTREXp/3xc-Myc as templates, respectively (primers 9 to 14; Table S1), that were later joined by overlap PCR using primers 9 and 14 (Table S1). Finally, the 2xTetO-MCS-3xc-Myc fragment was inserted into pTcINDEX-c-Myc linearized with XhoI using the NEBuilder^®^ HiFi DNA Assemble Master Mix to obtain the pTcINDEX/3xTetO/3xc-Myc vector.

### Inducible expression of *TcRDK2*

The PCR product (primers 1 and 2; Table S1) of the wild type *TcRDK2* gene (RDK2^WT^) was cloned into the pTcINDEX/3xTetO/3xc-Myc vector by MluI/EcoRV restriction sites. To generate a “kinase-dead” mutant, we mutated the conserved Lys70 (codon AAA) in the catalytic domain of TcRDK2 to alanine (codon GCA), producing the TcRDK2^K70A^ variant by overlap PCR (primers 1, 2, 15 and 16; Table S1), then the purified product was cloned into the pTcINDEX/3xTetO/3xc-Myc vector by the MluI/EcoRV restriction sites. Finally, a TcRDK2 variant lacking the last 97 C-terminal amino acids (TcRDK2^ΔPH^), corresponding to the predicted Pleckstrin Homology (PH) domain, was PCR-amplified (primers 1 and 16; Table S1) and cloned into the pTcINDEX/3xTetO/3xc-Myc vector via the MluI/XhoI restriction sites. Once obtained, these constructs were transfected into *T. cruzi* T7/TetR epimastigotes and culture for selection with hygromycin. We used 0.5 μg/mL tetracycline to induce expression in epimastigote cells and 1 μg/mL doxycycline for metacyclic trypomastigotes and infection essays.

### *TcRDK2* endogenous tagging

To perform CRISPR/Cas9-mediated endogenous C-terminal tagging of RDK2, the choice of the target sequence to design sgRNA templates was done as previously described (74–76). A donor DNA cassette containing the 3xc-Myc tag sequence and the puromycin resistance gene to induce HDR was amplified using the pMOTag23M vector (68) as template, and 60-nt long primers as previously described (30) (primers 19 and 20; Table S1).

For endogenous N-terminal tagging, the protospacer region was selected in the sense strand from nt -26 and -6 upstream the start codon of *TcRDK2*. To produce the donor DNA for N-terminal tagging we generated the pMOTagN23M plasmid to be used as DNA template in PCR. The pMOTagN23M plasmid was constructed from pMOTag2T by removing the Ty1 tag and inserting downstream the puromycin N-acetyltransferase (*PAC*) gene a copy of the *T. cruzi* HX1 trans-splicing sequence followed by three copies of the c-Myc tag (74). The donor cassette, amplified with 60-nt long primers containing 39 nt of each homologous region (HR) (primers 22 and 23; Table S1), contained HR1, *PAC*, 3xc-Myc tag sequence, and HR2 (Fig. S2).

Epimastigotes of a *T. cruzi* Y strain cell line that constitutively express both T7RNAP and Cas9 nuclease (T7RNAP/Cas9) were co-transfected with the sgRNA template and the donor DNA and then cultured for 3 weeks with G418 and puromycin for selection of resistant parasites. Endogenous gene tagging was verified by PCR from gDNA (primers 24 to 27; Table S1) and by western blot analysis using total protein extracts.

### *TcRDK2* ablation

We performed a CRISPR/Cas9-mediated knock out of *TcRDK2* using a standard strategy developed in our laboratory (30). Briefly, *T. cruzi* T7RNAP/Cas9 epimastigotes were transfected with a sgRNA template obtained by PCR (primers 28 and 29; Table S1) and two donor DNA cassettes amplified from p*BSD* and p*PAC* (primers 30 and 31; Table S1), respectively. The donor DNA was provided to induce HDR and contained a blasticidin or a puromycin resistance marker respectively flanked by 40 and 37-nt homologous regions corresponding to the 5’ and 3’ end of the *TcRDK2* UTRs. Selection of transfectants was done with puromycin and blasticidin to ensure both alleles were replaced by resistance markers. Gene knockout was verified by PCR on gDNA using a specific primer set (primers 25 and 26; Table S1). After *TcRDK2* knockout was confirmed, a clonal population was obtained by limiting dilutions.

### Cell transfections

*T. cruzi* Y strain epimastigotes were transfected as described previously (71). Briefly, *T. cruzi* epimastigotes in early exponential phase (4 x 10^7^ cells) were washed with PBS, pH 7.4, at room temperature (RT) and transfected in ice-cold CytoMix (120 mM KCl, 0.15 mM CaCl_2_, 10 mM K_2_HPO_4_, 25 mM HEPES, 2 mM EDTA, 5 mM MgCl_2_, pH 7.6) containing 25 μg of each plasmid construct and 25 μg of donor DNA in 4-mm electroporation cuvettes with three pulses (1500 volts, 25 microfarads) delivered by a Gene Pulser Xcell™ Electroporation System (Bio-Rad). Transfected epimastigotes were cultured in LIT medium supplemented with 20% heat-inactivated FBS until stable cell lines were obtained. When needed, the antibiotic used for drug selection and maintenance was added. Clonal populations of transfectant parasites were obtained by limiting dilutions in LIT medium and a final dilution in conditioned media (20% heat inactivated FBS, 40% filtered supernatant from WT cells in exponential phase, 40% LIT media, penicillin (100 I.U./mL), streptomycin (100 µg/mL), and appropriate antibiotics) to a final density of 2.5 cells/mL and plated 200 µl per well in 96-well plates.

### Western blot analyses

Western blots were performed as previously described (77). Briefly, parasites in exponential phase of growth were washed in 1x PBS pH 7.4 and resuspended in radio-immunoprecipitation assay (RIPA) buffer (150 mM NaCl, 20 mM Tris-HCl, pH 7.5, 1 mM EDTA, 1% SDS, 0.1% Triton X-100) plus a mammalian cell protease inhibitor cocktail (diluted 1:250), 1 mM phenylmethylsulfonyl fluoride, 2.5 mM tosyl phenylalanyl chloromethyl ketone, 100 M *N*- (*trans*-epoxysuccinyl)-L-leucine 4-guanidinobutylamide (E64), and Benzonase Nuclease (25 U/mL culture). After lysis the cells were then incubated for 30 min on ice, and protein concentration was determined by BCA protein assay. Thirty micrograms of protein from each cell lysate were mixed with 4x Laemmli sample buffer (Bio-Rad) supplemented with 10% β-mercaptoethanol, before loading into a 10% SDS–polyacrylamide gels. Electrophoresed proteins were then transferred onto nitrocellulose membranes with a Trans-Blot Turbo Transfer System (Bio-Rad). After transfer the membranes were stained with Ponceau red and an image was acquired for loading control using a ChemiDoc Imaging System (Bio-Rad). Membranes were then destained using PBS-T (PBS containing 0.1% Tween 20) and blocked with 5% nonfat dry milk in PBS-T overnight at 4°C. Then, the nitrocellulose membranes were incubated for 1 hour at room temperature, with the primary antibody: monoclonal anti-c-Myc (1:1000), monoclonal anti-tubulin (1:40,000). After three washes with PBS-T, blots were incubated with the secondary HRP-conjugated antibody (goat anti-mouse IgG, diluted 1:10,000). Membranes were washed three times with PBS-T and incubated with Pierce™ ECL Western Blotting Substrate (Thermo Fisher Scientific) in dark for 5 min. Lastly, images were acquired with a ChemiDoc Imaging System (Bio-Rad).

### Immunofluorescence analyses

*T. cruzi* parasites (epimastigotes, trypomastigotes or amastigotes) were washed with 1x PBS pH 7.4 and fixed with 4% paraformaldehyde (PFA) in 1x PBS pH 7.4 for 1 h at RT. Thereafter, cells were allowed to adhere to 1 mg/mL poly-L-lysine–coated coverslips and then permeabilized for 5 min with 0.1% Triton X-100. The coverslips were then washed 3 times with 1x PBS pH 7.4. Cells were then blocked with trypanosome blocking solution (3% bovine serum albumin (BSA), 1% fish gelatin, 5% normal goat serum and 50 mM NH_4_Cl, in PBS pH 7.4), overnight at 4°C. Next the cells were incubated with primary antibodies: monoclonal anti-c-Myc (1:100), diluted in 1% BSA in 1x PBS pH 8.0 for 1 h at RT. Cells were washed three times with 1% BSA in 1x PBS pH 8.0 and then incubated for 1 h at RT with secondary antibodies: Alexa Fluor 488–conjugated donkey-anti mouse (1:400). The incubation was performed keeping the cells protected from light to avoid photobleaching. Then, cells were washed 3 times with 1% BSA in 1x PBS pH 8.0 and mounted on slides using Fluoromount-G mounting medium containing 5 μg/mL 4,6-diamidino-2-phenylindole (DAPI) to stain genetic material. Differential interference contrast (DIC) and fluorescence optical images were captured using a Nikon Ni-E epifluorescence microscope on 100x oil immersion lens using NIS-Elements software (Nikon Instruments Inc), images were then deconvolved with the same software for 20 iterations using Richardson-Lucy method and automatic noise level. Images were processed with FIJI (ImageJ; National Institutes of Health, Bethesda, MD, USA).

### *In vitro* metacyclogenesis

Metacyclic trypomastigotes were obtained following the protocol described in (78) with some modifications. Briefly, 0.5 x 10^7^ *T. cruzi* epimastigotes in exponential phase were cultured for 4 days in LIT medium supplemented with 10% heat inactivated FBS. The parasites were then washed two times in 2 mL of triatome artificial urine (TAU) (190 mM NaCl, 17 mM KCl, 2 mM MgCl_2_, 2 mM CaCl_2_, 0.035% sodium bicarbonate, 8 mM phosphate, pH 6.9) and resuspended in 0.2 mL of TAU medium. Parasites were then incubated for 2 h at 28°C. After incubation, parasites were added to flasks and incubated horizontally for 96 h in 20 mL TAU 3AAG medium (TAU medium supplemented with 10 mM L-proline, 50 mM sodium L-glutamate, 2 mM sodium L-aspartate, and 10 mM glucose) in T25 flasks. For quantification of metacyclogenesis, the supernatant containing a mixture of epimastigotes, metacyclic trypomastigotes, and intermediate forms were centrifuged at 1300 x g for 15 min and fixed for 1h at RT in 4% PFA in PBS, attached to poly-L-Lysine-coated coverslips and washed three times with 1x PBS pH 7.4. Then, parasites were mounted onto glass slides with Fluoromount-G containing 10 μg/mL DAPI, for DNA staining. 20 fields/slide were analyzed on a Nikon epifluorescence microscope with a 100x objective under oil immersion in three independent experiments. Metacyclic trypomastigotes were distinguished from epimastigotes by the morphology and kinetoplast location in the cell body.

### *In vitro* infection assay

*T. cruzi* invasion and intracellular replication assays were performed using human foreskin fibroblasts (hFFs). First, 5 x 10^4^ hFFs in 1 mL of DMEM supplemented with 10% FBS were added to 12-well plates containing a sterile coverslip and allowed to attach overnight at 37°C with 5% CO_2_. The next day a swimming protocol was performed on tissue culture-derived trypomastigotes by centrifuging at 1700 x g for 15 min and incubating upright in a 50 mL conical tube for 4h at 37°C with 5% CO_2_. This allowed the competent trypomastigotes to swim out of the pellet into the supernatant. Next, the supernatant was spun down, and density of parasites was determined using a Neubauer chamber and resuspended to a concentration of 5×10^6^ parasites/mL. The hFFs in the 12-well plate were washed with DHANKS (Hank’s Balanced Salt Solution, Cytiva Marlborough, MA) and 1 mL of the parasite suspension (5×10^6^ parasites) was added for a multiplicity of infection (MOI) of 100 (100 trypomastigotes/cell). The infection was stopped after 4 h by washing the coverslips in the wells 5 times with DHANKS. Then 1 mL of DMEM with 2% FBS was added to slow down the proliferation of hFFs. Coverslips were removed from the plate after 24 h for the invasion assay, and after 72 h for the replication assay, and placed into a 12-well plate containing 4% paraformaldehyde in 1x PBS pH 7.4 for 1 h. The coverslips were then washed with 1x PBS pH 7.4 and mounted onto glass slides containing 15 μg/mL DAPI in Fluoromount G for DNA staining of parasites and mammalian cells. To quantify invasion, 20 fields/slide were visualized on a Nikon Ni-E epifluorescence microscope, and the number of infected and non-infected cells were counted. To quantify the replication of amastigotes, 60 infected host cells were visualized per assay on a Nikon Ni-E epifluorescence microscope and the number of amastigotes per infected cell were counted.

### Phosphoproteomic analysis

*Trypanosoma cruzi* epimastigotes in exponential phase of growth were incubated with 0.5 μg/mL of tetracycline for 48 h. Thereafter, 3 × 10^8^ cells were centrifuged at 1,000 × g for 15 min and washed twice with 5 mL of buffer A with glucose (BAG, 116 mM NaCl, 5.4 mM KCl, 0.8 mM MgSO_4_, 50 mM HEPES and 5.5 mM glucose, pH 7.3) at room temperature. The parasites were then resuspended in 1 mL of ice-cold lysis buffer (0.4% NP-40, 1 mM EDTA, 150 mM KCl, cOmplete™ Mini EDTA-free Protease Inhibitor Cocktail, Pierce Phosphatase inhibitor, 50 mM Tris-HCl, pH 7.5) and mixed at 4°C for 30 min with agitation on a rocking shaker. After lysis, the protein concentration of each sample was determined using BCA assay. Samples were sent to the University of Cincinnati Proteomics Laboratory (UCPL) (Cincinnati, OH). Samples were buffer exchanged 3x against a 1:1 aqueous dilution of the cell signaling urea lysis buffer containing phosphatase inhibitors. The urea lysis buffer contained 9 M urea, 20 mM hepes and 1mM b-glycerophosphate, 2.5 mM sodium pyrophosphate and 1 mM activated sodium orthovanadate. After the buffer exchange, the volumes were brought up to 1 mL with the urea lysis buffer. The samples were then probe sonicated for 3 cycles of 15 sec each at 15 W output. Insolubilized material was pelleted by a 20,000 × *g* and the samples were placed in a new tube. A 660 nm assay was performed on the samples and a volume containing 1.3mg was removed for each sample and brought up to 1 mL with 1x Urea lysis buffer. The protocol for digest preparation including reduction with DTT, alkylation with IAA and digestion with trypsin was according to the instructions from the Cell Signaling Technology^®^ (Cat. 5622). The samples were digested with modified trypsin (Worthington Biochemical, Cat. LS003740) for overnight at 37°C. The digested peptides were purified and concentrated by passing them over a C18 sep-pak cartridge (Waters™, Cat. WAT051910). The samples were then dried in a SpeedVac and resuspended in binding/equilibration buffer from the High Select™ TiO2 phosphopeptide enrichment kit (ThermoFisher Scientific, Cat. A39223) to be enriched using the TiO2 spin tips according to the manufacturer’s instructions.

The elution was immediately dried in a SpeedVac and resuspended in 0.1% Formic acid. 5 uL of each enriched sample was analyzed by nanoLC-MS/MS (Orbitrap Eclipse). The data was searched against a combined contaminant and a *T. cruzi* TcYC6 strain database release 66 (tritrypdb.org) using Proteome discoverer version 3.0 with the Sequest HT search algorithm (Thermo Scientific) and the LFQ quantitation workflow with the IMP-ptmRS and associated nodes added to calculate probabilities of phosphorylation at each site. Ratios were calculated in a pairwise manner and *P* values calculated using the *t*-test background method. Only phosphopeptides were used for calculating the ratios and for normalization. LFQ values detected exclusively in all replicates of a given condition (RDK2^WT^ or EV) were log_2_-transformed and column-wise Z-score–normalized using the median in Perseus (version 2.1.3.0) (Tables S2 to S7).

The upregulated phosphopeptides were required to have values in at least three RDK2^WT^ samples, whereas downregulated phosphopeptides were required to have values in at least two EV samples. Repeated peptide fragments were removed from the dataset. Gene Ontology annotations for phosphoproteins were retrieved from TriTrypDB (tritrypdb.org). For phosphorylation motif analysis, we used a non-redundant set of phosphopeptide sequences (unique sequence and modification site) derived from significantly regulated phosphosites and generated a library containing all identified phosphorylated sites with 5 amino acids upstream and downstream of the modified residue. This library was analyzed in MEME Suite 5.5.9 (meme-suite.org) using the Modification Motifs (MoMo) tool with the Motif X algorithm. Additional data visualizations were produced using GraphPad Prism v10 and RStudio with R 3.6.0+.

### Statistical analyses

Values are expressed as means ± Standard Deviation (SD). Statistically significant differences between treatments were compared using unpaired Student’s *t*-test, and two-way ANOVA tests with multiple comparisons, as mentioned in the legends of the figures. Differences were considered statistically significant for *P* < 0.05, and *n* refers to the number of independent experiments performed. All statistical analyses were performed using GraphPad Prism 9 (GraphPad Software, San Diego, CA).

## ACKNOWLEDGMENTS

Funding for this work was provided by the National Institute of Allergy and Infectious Diseases of the U.S. National Institutes of Health was supported by the U.S. National Institutes of Health (NIH Grant R21AI178573) and the American Heart Association (AHA Grants 23IPA1054779 and 26BIPA1622636). N.L. was supported by the U.S. National Institutes of Health (NIH Grants R00AI137322 and R21AI182544 to N.L.). The funding agencies had no role in the study design, data collection, and interpretation, or the decision to submit the work for publication. Opinions contained in this publication do not reflect the opinions of the funding agencies. We declare that we have no competing financial interests.

## Supplementary figure legends

**Figure S1.** TcRDK2 structural information. (A) Structural prediction of TcRDK2 using AlphaFold-3. Model confidence using per-residue confidence score (pLDDT) is indicated by color-code. N-terminus -N- and C-terminus -C- are indicated. (B) Predicted features of TcRDK2 displaying the ATP binding site, conserved elements and putative functional domains. The N-lobe and C-lobe of catalytic domain are shown in blue, while the activation loop and Pleckstrin homology (PH) domain are shown in yellow and red, respectively. In a close-up are indicated several relevant amino acid residues in the catalytic domain, as well as the HRD motif (His165/Arg166/Asp167), the Asp185 residue of the DFG motif, and the conserved threonine in the activation loop (Thr207). Model was made using AlphaFold-3.

**Figure S2.** Expression of endogenously N-terminal tagged 3xc-Myc*-TcRDK2*. (A) Schematic representation of CRISPR/Cas9 mediated strategy for the N-terminal tagging of *TcRDK2.* The Homologous-directed repair determines integration of 3xc-Myc and antibiotic resistance gene (*Pac*, Puromycin N-acetyltransferase) at 5’ end of *TcRDK2.* UTR, untranslated region; *Igr*, tubulin intergenic region; *HX1*, *T. cruzi* trans-splicing region. Horizontal *black arrows* indicate primers used for tagging verification. (B) PCR verification of integration of DNA donor in the 5’ end of the endogenous *TcRDK2* locus*. TcRDK2* allele size: WT, 325 bp; 3xc-Myc tagged, 1605 bp. Lanes: M, 1-kb plus ladder; T7/Cas9, parental control; cMyc RDK2, 3xc-Myc*-TcRDK2*; H_2_O, PCR negative control. (C) *TcRDK2* endogenous tagging was verified by western blotting using antibodies anti-c-Myc (c-Myc). Tubulin (Tub) was used as a loading control. T7/Cas9, parental control, cMyc RDK2, 3xc-Myc-*TcRDK2*. (E) Immunofluorescence analysis (IFA) showed localization of endogenously tagged 3xc-Myc-*TcRDK2* detected with anti-c-Myc antibodies (*green*) in *T. cruzi* epimastigotes. Merge of *green* signal (RDK2) and DAPI staining (*blue*), and DIC images are also shown. Scale bars = 5 μm.

**Figure S3.** Epimastigote growth rate of *TcRDK2* mutant cell lines. (A) Growth rate of control (T7/Cas9) and *TcRDK2*-KO (RDK2-KO) epimastigotes. Growth rate was estimated between days 3 to 7. Student’s *t*-test was applied to growth rates calculated from each growth curve (*n* = 3). Growth rates were normalized using T7/Cas9 as the control group. (B) Growth rate of EV (empty vector, control) and RDK2^K70A^ epimastigotes in the absence (Tet−) or presence (Tet+) of 0.5 μg/ml tetracycline. Growth rate was estimated between days 3 to 7. Two-way ANOVA with Tukey’s multiple comparisons test was applied to growth rates calculated from each growth curve (*n* = 3). Growth rates were normalized using the EV Tet− condition as the control group. ns, not significant.

**Figure S4.** Infection of hFF cells with *TcRDK2*-KO metacyclic trypomastigotes. hFF cells were incubated with T7/Cas9 (control) and *TcRDK2*-KO (RDK2-KO) metacyclic trypomastigotes. (A and B) After 6 days, the mammalian cells were full of T7/Cas9 amastigotes (upper panel), but only after 10 days was possible to observe a few cells with intracellular *TcRDK2*-KO undefined stage forms (bottom panel). (A) Brightfield microscope, 20x and 100x; (B) DAPI staining (*blue*) and DIC images are shown; scale bars = 20 μm. Black and white arrows indicate representative vacuoles in hFF cells containing RDK2-KO parasites.

**Figure S5.** Differentiation of TcRDK2^WT^, RDK2^ΔPH^ and TcRDK2^K70A^ epimastigotes in LIT medium. (A and B) Metacyclogenesis observed in (A) TcRDK2^WT^ and TcRDK2^K70A^, and (B) TcRDK2^K70A^ epimastigotes cultured in LIT medium after 120 hours of incubation with 0.5 μg/ml tetracycline. Cultures were started at 2.5 x 10^6^ parasites/ml. Epimastigotes, intermediate forms and metacyclic trypomastigotes were distinguished from epimastigotes by the morphology and kinetoplast location in the cell body. Data represent means, and error bars indicate S.D. from *n* = 3 independent experiments. \**P* < 0.05; \*\**P* < 0.01; \*\*\*\**P* < 0.0001. Statistical analysis was performed using two-way ANOVA with Sidak’s multiple comparisons test. For the results shown in panel A, only biologically relevant pairwise comparisons are indicated.

**Figure S6. Phosphoproteomic analysis of upregulated phosphosites: Heatmap of phosphosites uniquely detected in RDK2^WT^ samples.** Phosphopeptides identified exclusively in tetracycline-induced RDK2^WT^ cells (not detected in EV controls) were considered upregulated. For each phosphopeptide, normalized intensities across the three RDK2^WT^ replicates were converted to z-scores and sorted from highest to lowest mean z-score. Red indicates higher relative abundance within RDK2^WT^ samples, whereas blue indicates lower relative abundance.

**Figure S7. Phosphoproteomic analysis of upregulated phosphosites: Heat map of phosphosites uniquely detected in EV control samples.** Phosphopeptides identified exclusively in EV cells (absent from tetracycline-induced RDK2^WT^ samples) were considered downregulated upon RDK2^WT^ overexpression. For each phosphopeptide, normalized intensities across the three EV replicates were converted to z-scores and ordered from highest to lowest mean z-score. Red denotes higher relative abundance within EV samples, whereas blue denotes lower relative abundance.

